## Supplemental Figures for "Adult choline supplementation in a Down syndrome model reduces co-morbidities and improves cognition"

**Supplementary Figure 1 – Experimental diet composition.**

**A**

*Teklad Custom Diet*

**TD.140777**

**Choline Chloride Diet (1.1 g/kg, AIN-76A), G**

| Formula | g/Kg |
|---|---|
| Casein | 200.0 |
| DL-Methionine | 3.0 |
| Sucrose | 500.79 |
| Corn Starch | 150.0 |
| Corn Oil | 50.0 |
| Cellulose | 50.0 |
| Mineral Mix, AIN-76 (170915) | 35.0 |
| Vitamin Mix, AIN-76A (40077) | 10.0 |
| Choline Chloride | 1.1 |
| Ethoxyquin, antioxidant | 0.01 |
| Green Food Color | 0.1 |

**Footnote**

Modification of the AIN-76A formula (CA.170481) to include 1.1 g/kg choline chloride. Color coded green.

**Selected Nutrient Information[1]**

| | % by weight | % kcal from |
|---|---|---|
| **Protein** | 17.7 | 18.8 |
| **Carbohydrate** | 65.0 | 68.8 |
| **Fat** | 5.2 | 12.4 |
| **Kcal/g** | 3.8 | |

[1] Values are calculated from ingredient analysis or manufacturer data

**B**

*Teklad Custom Diet*

**TD.140778**

**Choline Chloride Diet (5 g/kg, AIN-76A), R**

| Formula | g/Kg |
|---|---|
| Casein | 200.0 |
| DL-Methionine | 3.0 |
| Sucrose | 496.89 |
| Corn Starch | 150.0 |
| Corn Oil | 50.0 |
| Cellulose | 50.0 |
| Mineral Mix, AIN-76 (170915) | 35.0 |
| Vitamin Mix, AIN-76A (40077) | 10.0 |
| Choline Chloride | 5.0 |
| Ethoxyquin, antioxidant | 0.01 |
| Red Food Color | 0.1 |

**Footnote**

Modification of the AIN-76A formula (CA.170481) to include 5 g/kg choline chloride. Color coded red.

**Selected Nutrient Information[1]**

| | % by weight | % kcal from |
|---|---|---|
| **Protein** | 17.7 | 18.8 |
| **Carbohydrate** | 64.6 | 68.7 |
| **Fat** | 5.2 | 12.4 |
| **Kcal/g** | 3.8 | |

[1] Values are calculated from ingredient analysis or manufacturer data

**A-B.** Experimental diets were modified AIN-76A chow from Envigo Teklad Diets. Other than the amount of choline chloride present, ChN diets (1.1 g/kg choline chloride; **A**) had the same general nutritional composition when compared to Ch+ diets (5 g/kg choline chloride; **B**).

**Supplementary Figure 2 – Radial arm water maze and IntelliCage testing.**

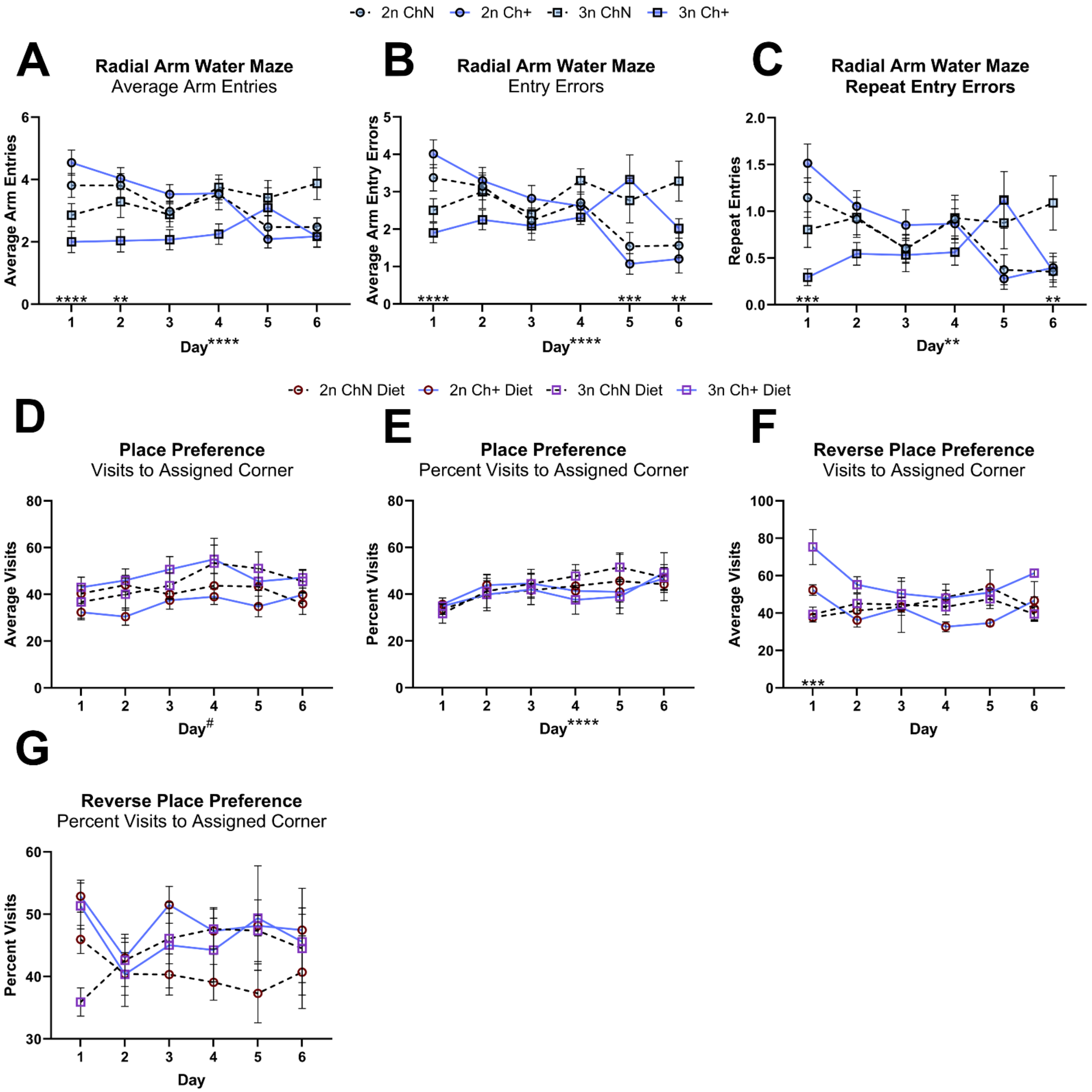


**A-C.** Animals were tested in the radial arm water maze (n = 5-9 per sex, diet and genotype) and assessed for the average number of arm entries (**A**), arm entry errors, defined as entries into an arm other than the goal arm, in trials where at least 1 arm was entered (**B**), and repeat entry errors, defined as re-entry into a previously entered arm in trials where at least 1 arm was entered (**C**). **D-G.** The IntelliCage apparatus was used (female mice only; n = 5-8 per diet and genotype) to examine performance in tasks where water access was limited to a single corner for each mouse, via both place preference (**D-E**) and place preference reversal tasks (**F-G**). We assessed the number of visits to the assigned corner (**D, F**) and percent of visits to the assigned corner (**E, G**).

Error bars represent SEM; *p $< 0.05$, **p $\leq 0.01$, ***p $\leq 0.001$, ****p $\leq 0.0005$ (MANOVA).

**Supplementary Figure 3 – Histological evaluation of steatosis in hepatic tissue.**

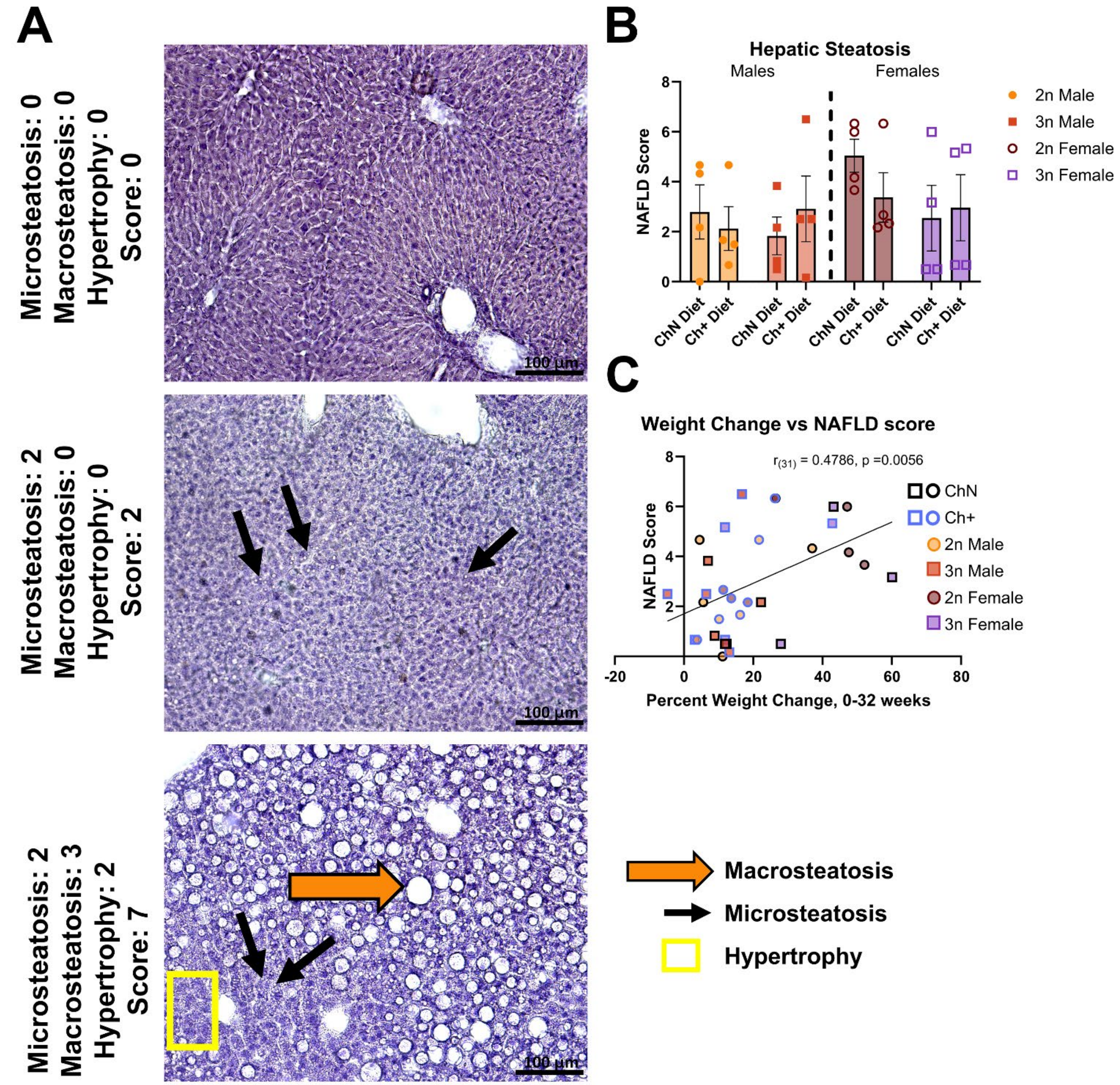


**A.** Representative low-magnification photomicrographs of hematoxylin- and eosin-stained hepatic tissue, demonstrating examples of scoring for non-alcoholic fatty liver disease (NAFLD; n = 4 animals per diet and genotype; sexes analyzed separately). The final score was the sum of scores for microvesicular steatosis, macrovesicular steatosis, and hepatocellular hypertrophy. Large orange arrows indicate macrosteatosis – fat droplets larger than the cell's nucleus, while small, black arrows indicate small liquid droplets indicative of microsteatosis. A yellow box indicates hepatic cells that show hypertrophy. Scale bar = 100µm. **B.** Average NAFLD scores across groups (MANOVA). **C.** NAFLD scores were correlated with percent weight change from 0-32 weeks on experimental diets (Pearson's correlation).
Error bars represent SEM.

**Supplementary Figure 4 – Additional endpoint peripheral cytokines elevated in 3n female mice.**

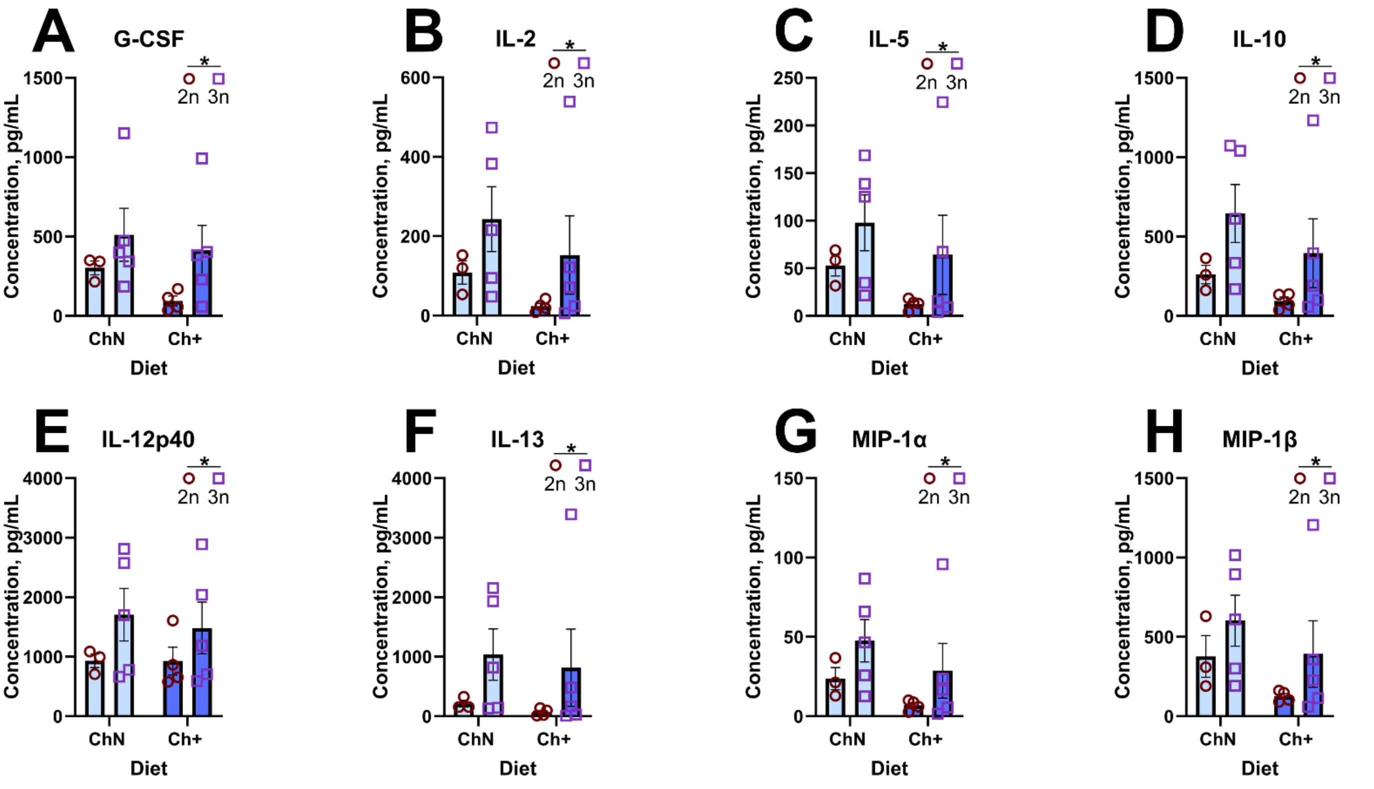


**A-G.** Analysis of peripheral plasma cytokines using the Bio-Plex® suspension array multiplexing system (n = 3-5 per diet and genotype). Individual cytokines that were significantly elevated in 3n female mice included granulocyte colony-stimulating factor (G-CSF; **A**), Interleukin (IL)-2 (**B**), IL-5 (**C**), IL-10 (**D**), IL-12p40 (**E**), IL-13 (**F**), Macrophage inflammatory protein (MIP)-1α (**G**), and MIP-1β (**H**).
Error bars represent SEM; *p < 0.05 (MANOVA).
